# The bioactive dietary polyphenol preparation alleviates depression and anxiety-like behaviors by modulating the regional heterogeneity of microglia morphology

**DOI:** 10.1101/2023.03.30.534961

**Authors:** Eun-Jeong Yang, Tal Frolinger, Susan Westfall, Umar Haris Iqbal, James Murrough, Giulio M. Pasinetti

## Abstract

**Scope:** The goal of this study is to investigate the effects of a bioactive dietary polyphenol preparation (BDPP), which is made up of grape-derived polyphenols, on microglial responses, as well as the underlying molecular mechanisms in depression and anxiety-like behaviors.

**Methods and results:** We find that treatment with BDPP significantly decreased depression-like and anxiety-like behaviors induced by chronic stress in mice, while leaving their locomotor activity unaffected. We also find that BDPP treatment reversed microglia activation in the amygdala and hippocampal formation, regions of the brain involved in emotional regulation, from an amoeboid shape to ramified shape. Additionally, BDPP treatment modulates the release of pro-inflammatory cytokines such as interleukin-6 via high mobility box 1 protein and the receptor for advanced glycation end products (HMGB1-RAGE) signaling pathway in activated microglia induced by chronic stress.

**Conclusion:** Our findings suggest regional heterogeneity in microglial responses following chronic stress in subregions of the corticolimbic circuit. Specifically, activation of the immune-inflammatory HMGB1-RAGE pathway might provide a new avenue for therapeutic intervention in stress-induced anxiety- and depression-like behavior, using bioactive and bioavailable polyphenols.

## 1. INTRODUCTION

Dietary polyphenols have emerged as a promising therapeutic approach for brain disorders such as major depressive disorder (MDD), with studies indicating that increasing intake of polyphenol-rich foods or supplements may improve mood and attenuation of MDD behavior [1, 2]. However, it is important to note that the therapeutic effectiveness of individual polyphenols, including resveratrol (RSV), flavonoids, and anthocyanins, may vary in the context of MDD, depending on factors such as their composition, dosage, and duration of treatments [3]. Further research is necessary to fully understand the underlying mechanisms in the brain that mediate the effects of polyphenols in MDD.

The therapeutic benefits of dietary polyphenols may be due to the presence of complex mixtures of bioactive compounds, which can have greater efficacy than single purified compounds by acting on multiple targets and producing synergistic effects [4]. We recently developed bioactive dietary polyphenol preparation (BDPP) which is a combination of three bioactive and commercially available polyphenol products, particularly a grape seed polyphenol preparation (GSPE), concord grape juice (CGJ), and RSV [5, 6]. This preparation is designed to simultaneously target multiple pathogenic mechanisms [7]. For instance, it has been reported that BDPP derived bioactive metabolites have the ability to penetrate the blood-brain barrier, which is an important step towards its potential use as a dietary supplement for brain health [8]. We also identified that BDPP promoted psychological resilience against stress-induced depressive-like behavior by modulating neuronal synaptic plasticity and peripheral pro-inflammatory cytokines [9].

Activation of microglia due to psychological and environmental stress is considered a major contributor to the development of MDD [10]. This activation triggers immune responses that release pro-inflammatory cytokines, including IL-1β and IL-6 which can lead to changes in the structure and function of neurons, ultimately resulting in development of MDD [11-13]. Recent studies have revealed that microglia exhibit diverse cellular and molecular features at baseline, indicating the existence of distinct subpopulations and subtypes that differ in their functions [14]. It is important to take into account the heterogeneity of microglial responses in investigating the underlying mechanisms in MDD development and treatment, which could lead to the development of more effective therapies.

Although BDPP has demonstrated its potential in regulating mood and decreasing the likelihood of depression and anxiety, its mechanisms of action on the complex relationship between microglial regional heterogeneity and inflammatory responses in the brain and neuropsychiatric impairments are not yet fully understood. The aim of the present study is to investigate the molecular mechanisms behind the impact of BDPP on the heterogeneous immune responses of microglia in different brain regions, ultimately clarifying potential mechanisms associated with polyphenolic treatments and mitigation of MDD.

## 2. MATERIALS & METHODS

### 2.1 Chemicals

Polyphenol-free diet (AIN-93G) was purchased from Research Diets, Inc. (New Brunswick, NJ). Food-grade resveratrol was purchased from ChromaDex (Irvine, CA, USA). GSPE was purchased from Supplement Warehouse (UPC 603573579173, Bolingbrook, IL, USA). Concord purple grape juice (Welch Foods Inc., Concord, MA, USA). One lot of the resveratrol and one lot of the GSPE were used for this particular study and were stored at 4 °C in the dark.

### 2.2 Animals

Cx3Cr1^CreErt2/+^ and Eef1a1^LSLeGFP10a/+^ male mice were purchased from the Jackson Laboratory. Eef1a1^LSLeGFP10a/+^ were crossed with Cx3Cr1^CreErt2/+^ mice to generate microglia glia-specific TRAP mice. All male mice were maintained on a 12:12-h light/dark cycle with lights on at 07:00 h in a temperature-controlled (20 ± 2 °C) in the centralized animal care facility of the Center for Comparative Medicine and Surgery at the Icahn School of Medicine at Mount Sinai. Mice were fed a polyphenol free diet (AIN-93G). All procedures were approved by the Mount Sinai Institutional Animal Care and Use Committee (IACUC-PROTO202000019) and conducted in accordance with the guidelines and regulations.

### 2.3 Stress exposure and BDPP treatment

For assessing BDPP’s effect in a model of chronic stress (**Figure 1b**), following 2-weeks of polyphenol-free diet, 2 months of age mice were randomly subdivided for treatment groups including: non-stressed mice receiving regular drinking water (n=6, unstressed) and stressed mice exposed to chronic stress for 28 days (n=13, stressed). Animals were subjected to chronic stress exposure that consisted of a random combination of two stressors per day (**Supplemental Table 1**). Mice in the stressed groups were treated with regular drinking water (n=6) or with BDPP prior to the stress exposure (n=7). BDPP, composed of GSPE, RSV and CGJ, was delivered through the drinking water. The total amount of polyphenol delivered was based on a calculation of human consumption of polyphenols estimated at 1 g/day, with phenolic acids accounting for approximately 30% of the total intake, or 330 mg/day [15]. To determine the equivalent dose in mice based on body surface, a USDA recommended formula was used, revealing that a human dose of 330 mg/day total polyphenol corresponds to 62 mg/kg/day in mice. In this study, the daily intake of GSE was calculated to be 200 mg/kg, RSV was 400 mg/kg, and the total polyphenols from juice extract were 183 mg/kg, which is approximately equivalent to 62 mg/kg/day in mice [8, 9, 16-18]. These doses were also chosen based on the equivalent doses used in a previous study that showed efficacy in animal models for each component(citation). For all experiments, body weight was assessed weekly.

**Figure 1.**
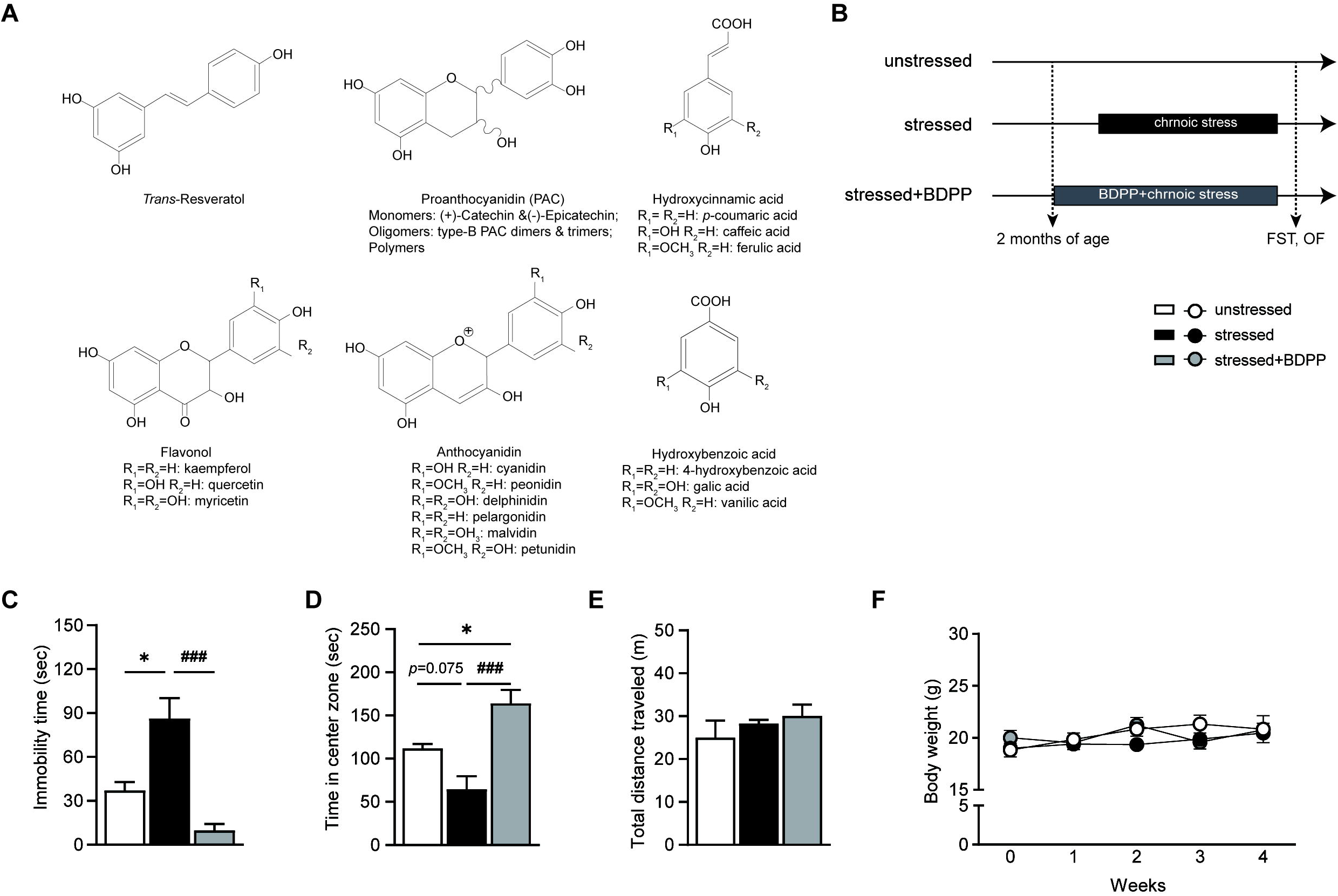
BDPP administration protects against chronic stress induced depression and anxiety-like behaviors. (A) Representative structures of polyphenols (Trans-resveratol, PAC, hydroxycinamic acid, flavonol, anthocyanidin, hydroxybenzoic acid) in BDPP. (B) Experimental outline. The experimental design consisted of a control group of unstressed mice, an age-matched group of mice and two groups of stressed mice that were either treated with regular drinking water or with BDPP. Open field test (OF) and forced swimming test (FST) were assessed after exposure to chronic stress. (C) A bar graph showing immobility time in the FST. (D-E) Bar graphs showing time in the center zone (D) and total distance traveled (E) in OF. (F) A line graph showing the average body weight over time for subjects undergoing chronic stress and undergoing chronic stress with treatment. One-way ANOVA was followed by Tukey Multiple comparison post hoc test, (n=6-7, \**p* < 0.05, compared to unstressed mice; ^###^*p* < 0.001, compared to stressed mice). Data are presented as mean ± SEM.

### 2.4 Open field test (OF)

Mice were placed into the center of box (40×40×40 cm) in a brightly lit room (150 lux). During the 10-min session, mice were scored for the total distance traveled and time spent in the center of the box. Animal behavior was video-recorded, tracked, and analyzed with ANY-maze™ tracking software (Stoelting Co., IL, USA).

### 2.5 Forced swimming test (FST)

Mice were individually put into 15-cm-diameter glass cylinder filled to 20 cm with 23□±□1 °C water. Immobility, defined as the absence of all movement except motions required maintaining the animal’s head above the water, was independently evaluated from video clips during a 6-min session. Results were expressed as time (in seconds) that animals spent immobile during the final 5-min of the session. The behavior of the animals in all of the tests was recorded on video, and then tracked and analyzed using ANY-maze™ tracking software.

### 2.6 Immunofluorescence

Mice were anesthetized with ketamine (120 mg/kg) and xylazine (24 mg/kg) and perfused transcardially with 10 mL PBS. Brains were then removed, fixed in formalin, and incubated in 0.02% sodium azide for bacterial contamination prevention. Brains were cut using a microtome in 5-μ m sections, and were stored at 4 °C. Brain sections were washed with PBS and permeabilized with PBS + 0.2% Triton X-100 (PBST) followed by blocking with 2% normal goat serum in PBST. Brain sections were incubated with primary antibody (Iba-1, 1:1000, Wako Chemicals, VA, USA) in 2% normal goat serum in PBST overnight at 4 °C. Brain sections were then washed and incubated with Alexa Fluor-conjugated secondary antibody (Alexa Fluor 568-labeled goat anti-rabbit, Thermo Fisher Scientific, MA, USA) at 1:500 in 2% normal goat serum in PBST for 1 h at RT. Brain sections were washed and coverslipped using Prolong Gold anti-fade with DAPI (Thermo Fisher Scientific) and dried overnight.

### 2.7 Morphological analysis of microglia

Imaging was performed using a Zeiss LSM 780 Confocal Microscope (Zeiss, DE, Germany) for collapsed 2 dimensional (2D) images and 2D or 3D image processing was performed using Zen software (Zeiss) or IMARIS 9.1.2 (Bit Plane Inc, MA, USA), respectively. Neurolucida Explorer (MBF Bioscience, VT, USA) was used to manually trace and generate analyses of microglia from the amygdala (AG), hippocampal formation (HF) and medial prefrontal cortex (mPFC) regions. One microglial cell was selected per 0.045 μ m^2^ of each images. The cell body or soma of microglia was traced, and the number of branch tips were measured. Sholl analyses were performed for each cell to account for complexity of microglia. Concentric circles were spaced 10 μm apart originating from the soma, and intersections of each cell with concentric circles were measured. We used n=3 per groups for brain region comparisons.

### 2.8 Purification of microglia by fluorescence-activated cell sorting

Microglia from one brain hemisphere were isolated as previously described [19]. In brief, half hemispheres were digested in a 0.4 mg/ml collagenase D solution (Roche Diagnostics, Germany) in phenol-free RPMI media supplemented with 10 mM HEPES and 5% FBS for 30 min at 37°C, homogenized using a 18G syringe in Hank’s Balanced Salt Solution without calcium or magnesium ions (Thermo Fisher Scientific) and passage through a 70-mm nylon cell strainer. Homogenates were centrifuged at 500 g for 5 minutes. Supernatants were removed and cell pellets were re-suspended in 37% isotonic Percoll (GE Healthcare, IL, USA) at room temperature. A discontinuous Percoll density gradient was layered as follows: 70%, 37%, 30%, and 0% isotonic Percoll. The gradient was centrifuged for 40 minutes at 500 g without break, and microglia were collected from the interphase between the 70% and 37% Percoll layers. Each hemisphere extraction yielded ∼ 2×10^3^ viable cells and 90% microglia. Cells were washed and then incubated with FC Block (1:1000, BD biosciences, CA, USA) for 30 min on ice. Cells were then incubated with the conjugated antibodies CD45-PerCP (1:1000, BD biosciences), CD11b-PE Cy7 (1:1000, BD biosciences) for 1 hr on ice. Cells were washed and then resuspended in FACS buffer (2% fetal bovine serum in PBS with 1 mg/mL sodium azide) for analysis. Nonspecific binding was assessed by using nonspecific, isotype-matched antibodies. Antigen expression was determined using the BD LSRII FACS analyzer (BD biosciences). A total of 20,000 events were recorded for each sample and isotype matched conjugate. Data were analyzed using BD FACSDiVa software and gating for each antibody was determined based on nonspecific binding of appropriate negative isotype–stained control samples.

### 2.9 Extraction of RNA from isolated microglia cells

Microglial RNA was purified using the Dynabeads® mRNA DIRECT™ Micro Kit (Thermo Fisher Scientific). RNA concentration was determined by NanoDrop spectrophotometry (PeqLab Biotechnology, Erlangen, Germany). The optical density ratio of 260/280 was measured using Nanodrop spectrophotometer (PeqLab Biotechnology) and ranged between 1.9 and 2.1. RNA samples were stored at −80°C.

### 2.10 Real-Time Quantitative Reverse Transcription PCR (qRT-PCR)

Total of 200 ng RNA from isolated microglia cells was used to synthesize complementary DNA using the SuperScript VILO cDNA Synthesis Kit (Invitrogen, CA, USA, 11754050). Gene-specific primers for *Hmbg1, Rage, IL-1*β and *IL-6* were designed using Primer blast (**Supplemental Table 2**), and gene expression was measured in 4 replicates by qRT-PCR using Maxima SYBR Green master mix (Thermo Fisher Scientific). Hypoxanthine phosphoribosyltransferase (*HPRT*) expression level was used as an internal control. Data were normalized using the 2^-^ΔΔ^Ct^ method. Levels of target gene mRNAs were expressed relative to those found in unstressed mice and plotted in GraphPad Prism 9 software (CA, USA).

### 2.11 Statistical Analysis

All values are expressed as mean and standard error of the mean (SEM). For statistical analysis of behavioral tests as well as biochemical analyses, Multiple t-test or One-way ANOVA followed by turkey’s test were used. In all studies, outliers were excluded and the null hypothesis was rejected at the 0.05 level. All statistical analyses were performed using GraphPad Prism 9 software.

## 3. RESULTS

### 3.1 BDPP treatment moderates susceptibility to chronic stress-induced depression and anxiety-like behaviors

As shown **Figure 1A**, BDPP contains various grape-derived polyphenols, including GSPE, CGJ, and RSV. Both GSPE and CGJ are complex sources of dietary polyphenols with multiple chemical components such as proanthocyanidins (PAC), anthocyanidins, flavonols, and phenolic acids.

To examine the effects of BDPP on chronic stress-induced behavior deficits, mice were assigned to one of three groups: an unstressed control group (unstressed), a group subjected to chronic stress (stressed), and a group pre-treated with BDPP prior to exposure to chronic stress (stressed+BDPP) **(Figure 1B)**. Depression-like behaviors were evaluated using FST **(Figure 1C)** and anxiety-like behaviors were assessed using the OF **(Figure 1D and 1E)**. We found that 28-days of chronic stress resulted in depression-like behavior, as indicated by a significant increase in immobility time in FST **(Figure 1C, \**p*<0.05)** and anxiety-like behavior, as assessed by a decreased tendency to spend time in the center of the open field arena compared to unstressed control mice **(Figure 1D, *p*=0.075)**. Interestingly, BDPP administration significantly attenuates both depression-like and anxiety-like behaviors **(Figure 1C and 1D**, ^**###**^***p*<0.001)**. Mice exposed to chronic stress or chronic stress with BDPP treatment did not show any significant changes in locomotor activity **(Figure 1E)** or body weight **(Figure 1F)** compared to unstressed control mice.

This finding suggests that BDPP administration attenuates depression-like and anxiety-like behaviors induced by chronic stress without affecting locomotor activity or body weight.

### 3.2 BDPP treatment reduces microglial activation induced by chronic stress in AG and HF

Previous studies have demonstrated that chronic stress-induced alterations in microglial morphology in brain regions including the AG, HF, and mPFC can lead to a heterogeneous, pro-inflammatory state, supporting the existence of distinct subpopulations with potentially different functions under certain brain injury conditions [20].

To assess the effects of chronic stress and BDPP treatment on microglial morphological phenotypes, such as alterations in soma size and branch length in AG, HF, and mPFC, we conducted a Sholl analysis by 3D reconstruction of Iba-1 immunopositive microglia cells **(Figure 2A, 2D and 2G)**. We observed no significant differences in soma size among all groups in the AG, HF, and mPFC **(Figure 2B, 2E and 2H)**.

**Figure 2.**
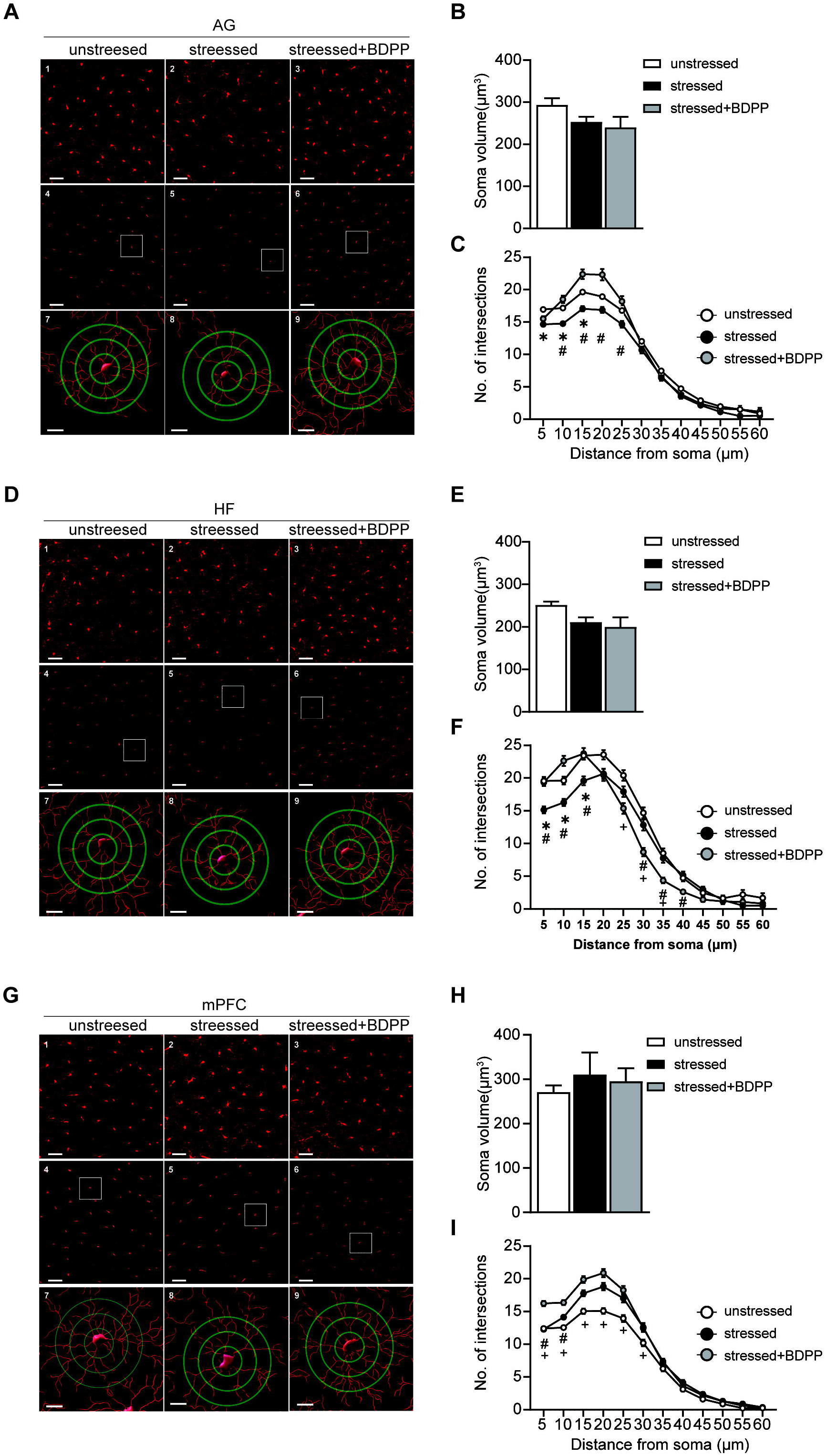
BDPP administration protects against the morphological effects on microglia induced by chronic stress in various brain regions. (A, D and G) The upper panels showing representative immunohistochemical images depicting Iba-1 immunopositive cells within the amygdala (A, AG), hippocampal formation (D, HF), and medial prefrontal cortex (G, mPFC). The middle panels showing 3D reconstructions of Iba-1 immunopositive cells, and in the lower panels, the branches of each microglia cell were measured by increasing radii in 10 µm steps to determine branch intersections. The area outlined with a white box in the middle panel is magnified in the lower panels. Length bars, 50 μm and 10 μm in upper/middle and lower panels, respectively. (B-C, E-F and H-I) Bar graphs showing soma size (B, E and H) and number of intersections (C,F and I) of 3D reconstructed microglia within the amygdala, hippocampal formation and mPFC, respectively. Multiple t-test was followed (n = 3 per group, 50 cells per group; \**p* < 0.05, compared to unstressed mice; ^#^*p* < 0.05, compared to stressed mice; ^+^*p* < 0.05, compared to stressed+BDPP mice). Data are presented as mean ± SEM.

However, stressed mice exhibited a significant reduction in the number of intersecting segments further away from the soma (5-15 µm away from the soma) in the AG **(Figure 2A panel 7 vs panel 8 and Figure 2C, \**p*<0.05)** and HF **(Figure 2D panel 7 vs panel 8 and Figure 2F, \**p*<0.05**), indicating a state of activation with decreased branch complexity. Importantly, BDPP treatment significantly restored the decreased branch complexity caused by chronic stress in the AG at 10-25 µm away from the soma **(Figure 2A panel 8 vs panel 9 and Figure 2C**, ^**#**^***p*<0.05)** and in the HF at 5-15 µm and 30-40 µm away from the soma **(Figure 2D panel 8 vs panel 9 and Figure 2F**, ^**#**^***p*<0.05)**.

Additionally, we did not observe any significant changes in microglial morphology in the mPFC following chronic stress compared to unstressed mice **(Figure 2G panel 7 vs panel 8 and Figure 2I)**. However, there were significant BDPP effects observed at 10-25 µm away from the soma compared to unstressed mice **(Figure 2G panel 9 vs panel 7 and Figure 2I**, ^**#**^**p<0.05)**.

The finding that chronic stress induces a region-specific microglial pro-inflammatory response, characterized by decreased complexity ramification in the AG and HF suggests the existence of distinct subpopulations. Most importantly, our finding demonstrates that BDPP treatment can mitigate these effects, offering insights into the potential mechanisms underlying its therapeutic effects on depression and anxiety-like behaviors via modulation of immune inflammatory responses.

### 3.3 BDPP treatment modulates pro-inflammatory response in activated microglia induced by chronic stress

HMGB1, one of damage-associated molecular patterns (DAMP), is a crucial mediator of immune inflammatory responses through binding RAGE receptor in activated microglia in response to stress [21]. To explore the molecular mechanisms underlying the therapeutic effects of BDPP on activated microglia, we investigated transcriptional changes in enriched microglia. To enable further analysis, microglial cells were isolated from the mouse brain through FACS, using CD11b and CD45 antibodies **(Figure 3A)**.

**Figure 3.**
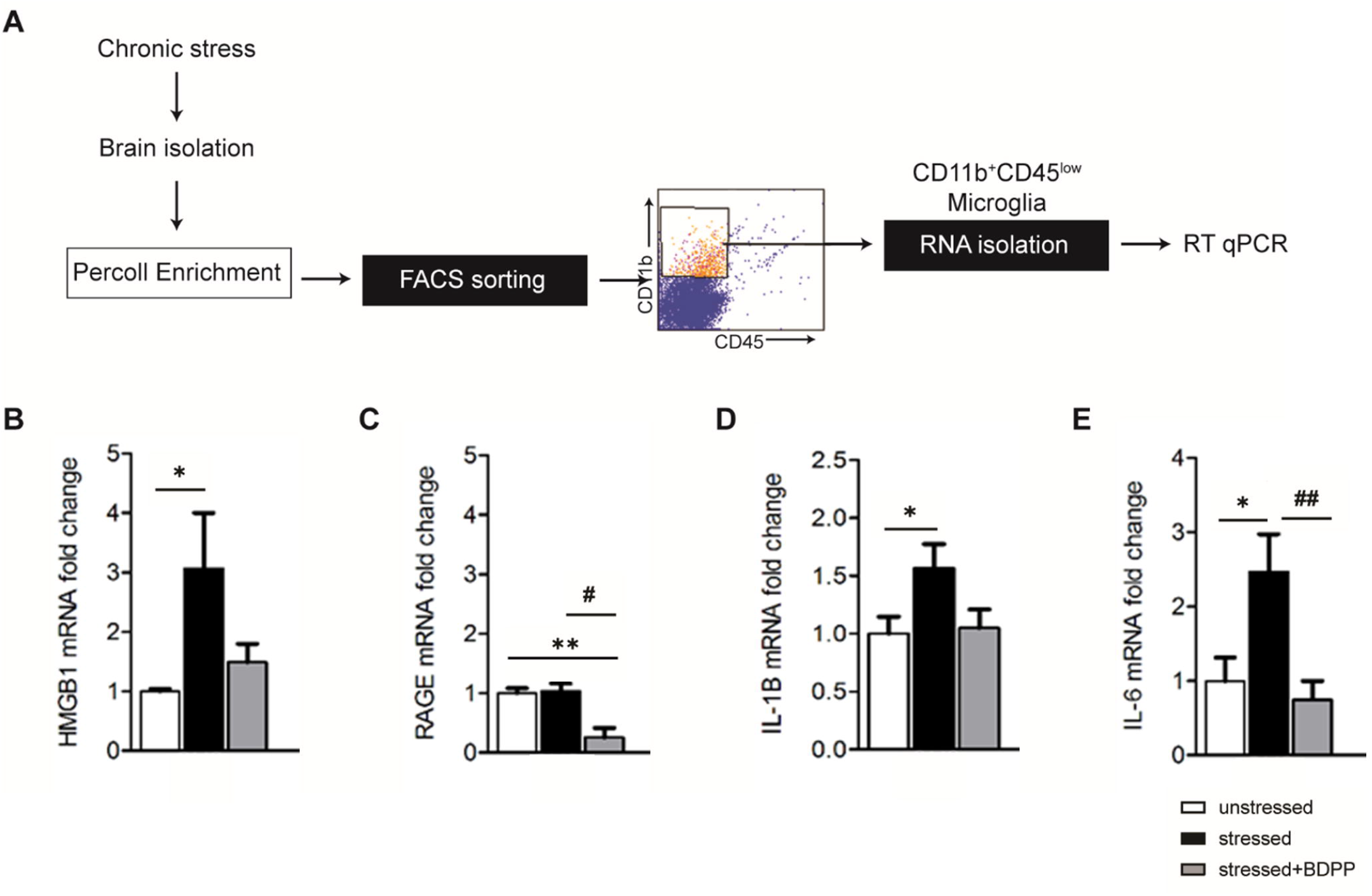
BDPP administration normalized upregulation of HMBG1-RAGE signaling pathway induced by chronic stress in enriched microglia. (A) Experimental paradigm for flow cytometry separation of microglia cells from brain. (B-E) Bar graphs showing relative quantification of mRNA expression of *Hmbg1* (B), *Rage* (C), I*l-1*β (D) and *Il-6* (E) in enriched microglia. One-way ANOVA was followed by Tukey Multiple comparison post hoc test, (n=6-7, \**p* < 0.05, \*\**p* < 0.01, compared to unstressed mice; ^#^*p* < 0.05, ^##^*p* < 0.01, compared to stressed mice). Data are presented as mean ± SEM.

We observed a significant 3-fold increase in HMGB1 mRNA expression in microglia enriched from stressed mice compared to unstressed control mice and found that BDPP treatment mitigated the HMGB1 mRNA expression in stressed group **(Figure 3B, \**p*<0.05)**. We found no significant increase in RAGE mRNA expression following exposure to chronic stress, when compared to unstressed mice. However, BDPP treatment significantly decreased RAGE mRNA expression compared to both unstressed and stressed mice **(Figure 3C, \*\**p*<0.01**, ^**#**^***p*<0.05)**. Our results indicate that chronic stress induces microglia activation through HMGB1-RAGE signaling pathway which has been linked to increased levels of microglia-driven pro-inflammatory cytokines, including IL-1β and IL-6.

We find that stressed mice exhibited a 1.5 and 2.5-fold increase in mRNA expression of the pro-inflammatory cytokines, IL-1β and IL-6, respectively **(Figure 3D and 3E, \**p*<0.05)**. BDPP treatment mitigated the IL-1β mRNA expression in the stressed group **(Figure 3D)**, while it significantly decreased IL-6 mRNA expression to the level observed in unstressed control mice **(Figure 3E**, ^**##**^ ***p* <0.01)**.

Together, our results suggest that chronic stress induces microglial activation through the HMGB1-RAGE signaling pathway, leading to increased levels of pro-inflammatory cytokines, and that treatment with BDPP can attenuate these inflammatory responses, potentially contributing to the mitigation of depression and anxiety-like behaviors.

## 4. DISCUSSION

Our study provides further evidence that bioavailable dietary polyphenolic preparations can modulate microglia-mediated immune inflammatory responses and mitigate anxiety- and depression-like behaviors, within the context of the regional heterogeneity of microglial responses to chronic stress in the brain, consistent with previous findings [22-24].

BDPP, a dietary supplement comprised by the grape-derived botanical supplements, GSPE, CGJ, and RSV [5, 6], is widely recognized for its anti-inflammatory properties as reflected in several disease models [2, 25]. Although there is a growing amount of evidence indicating that polyphenols have beneficial effects, the molecular mechanisms through which they exert their actions especially in brain have yet to be fully elucidated. Most importantly, the ability of BDPP in generating bioavailable bioactive polyphenols through dietary supplementation has allowed for several investigations, particularly related to brain disorders, due to the evidence showing bioactive metabolites reaching the brain at physiological level [5, 6, 8, 9]. Building upon this, the current study aims to investigate the potential role of BDPP in targeting the anti-inflammatory responses and underlying mechanisms associated to the mitigation of the depression and anxiety-like behaviors.

Our results tentatively suggest that BDPP treatment attenuates chronic stress-induced anxiety- and depression-like behaviors, possibly through modulation of microglial responses **(Figure 1 and 2)**. The morphology of microglia is strongly linked to their functional state, with activated microglia exhibiting an amoeboid shape in response to stress conditions, which can ultimately impair neuronal function through excessive synaptic pruning [26]. For example, it was found that BDPP influences the expression of molecules such as cAMP response element-binding protein (CREB) or c-Fos in AG, which are known for their role in the mechanisms of synaptic plasticity and neuronal function [27]. Further research is necessary to comprehensively investigate the effects of dietary polyphenols including bioactive metabolites able to restore microglial-neurons cross talking involving depression- and anxiety-like behaviors.

The corticolimbic circuit, which includes the AG, HF, and mPFC, is known to play a critical role in chronic stress-induced psychological impairments, possibly through the modulation of microglia function in this circuit. [27]. Interestingly, our study found that BDPP had varying effects on microglial morphology in different corticolimbic regions, with a restoration of microglial branching or complexity observed in the amygdala and hippocampal formation, but not in the prefrontal cortex **(Figure 2)**. Recent evidence has shown that microglia exhibit heterogeneity in different regions of the CNS under normal conditions, as demonstrated by various techniques including single-cell mass spectrometry and single-cell RNA sequencing [14, 28, 29]. Because of this, it is not unexpected that distinct subtypes of microglia, as we found in our study in the corticolimbic region, would differentially respond to stress and pharmacological intervention. Consistent with this hypothesis, we found that BDPP differentially influenced microglial morphology and transcription levels in different subregions of the corticolimbic circuit, which ultimately might have contributed to the mitigation of behavioral deficits.

Chronic stress has been largely shown to induce an increased expression of pro-inflammatory mediators in activated microglia [30]. For example, HMGB1 can be actively secreted from activated microglia or passively released from necrotic cells to initiate inflammatory responses under stress conditions [31]. Extracellular HMGB1 interacts with several pathogen recognition receptors such as the RAGE, to promote the local generation of pro-inflammatory cytokines including tumor-necrosis factor and IL-6 in brain [32]. HMGB1 binding to RAGE can also increase RAGE expression and cathepsin B activity, which triggers NLRP3 inflammasome activation and caspase-1-mediated pyroptosis with the release of IL-1β [33-35]. We find that chronic stress exposure increased the mRNA levels of HMGB1 and RAGE, as well as the expression of pro-inflammatory cytokines like IL-1β and IL-6, in enriched microglia. Interestingly, treatment with BDPP was able to mitigate HMGB1 and RAGE mRNA expression in enriched microglia, coinciding with attenuation of IL-1β and IL-6 **(Figure 3)**. This evidence tentatively suggests that BDPP might be effective in mitigating the onset of depressive and anxiety-like behaviors in stressed mice through the microglial HMGB1-RAGE signaling pathway.

To our knowledge, this study provides the first evidence supporting the hypothesis that dietary polyphenols may attenuate stress-induced depression- and anxiety-like behaviors, potentially through a mechanism associated with differential microglial activation in the corticolimbic pathways. Our findings suggest that there is regional heterogeneity in microglial responses following chronic stress in subregions of the corticolimbic circuit, specifically through the activation of the immune inflammatory HMGB1-RAGE pathway. This may provide a new avenue for therapeutic intervention in stress-induced anxiety- and depression-like behavior with bioactive and bioavailable polyphenols

## Supporting information

Supplemental Table 1, and will be used for the link to the file on the preprint site.

Supplemental Table 2, and will be used for the link to the file on the preprint site.

## FIGURE LEGENDS

**Supplemental Table 1**. Chronic stress schedule

**Supplemental Table 2**. Primers used in this study.

## Acknowledgements

This study was supported by Grant Number U19-AT010835 from the NCCIH and the ODS, and by the VA MERIT grant BX005054. Dr. Pasinetti holds a Senior VA Career Scientist Award. We acknowledge that the contents of this study do not represent the views of the NCCIH, the ODS, the NIH, the U.S. Department of Veterans Affairs, or the United States Government.

## Author Contributions

The authors’ contributions were as follows: E.Y, T.F, S.W, and UI participated in the acquisition and/or analysis of data. E.Y, J.M and G.M.P participated in the design and/or interpretation of the reported experiments or results. E.Y and G.M.P wrote the manuscript and all other authors reviewed and edited the manuscript

