## Supplementary figures and images for "The bioactive dietary polyphenol preparation alleviates depression and anxiety-like behaviors by modulating the regional heterogeneity of microglia morphology"

### Supplemental Table 1, and will be used for the link to the file on the preprint site.

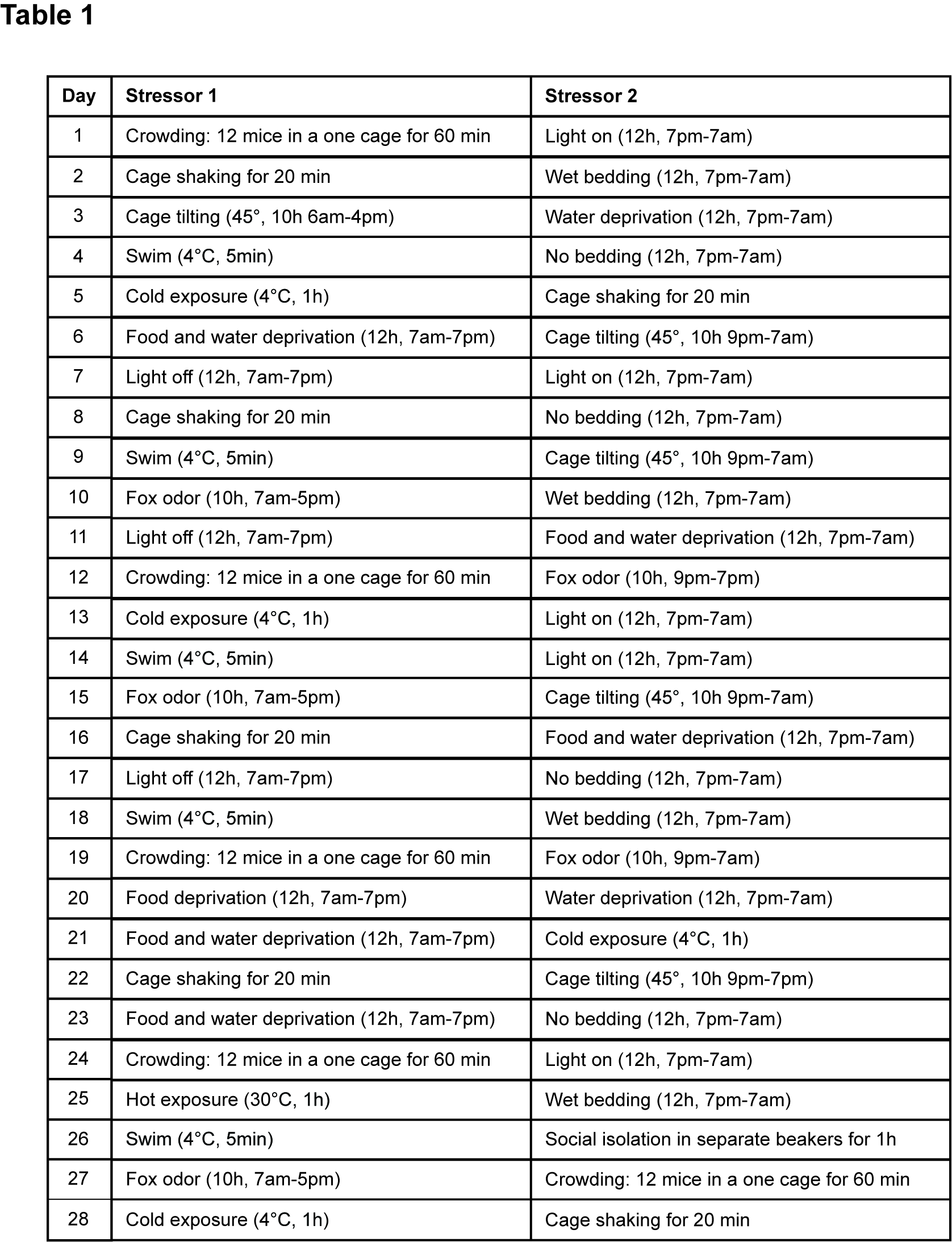

### Supplemental Table 2, and will be used for the link to the file on the preprint site.

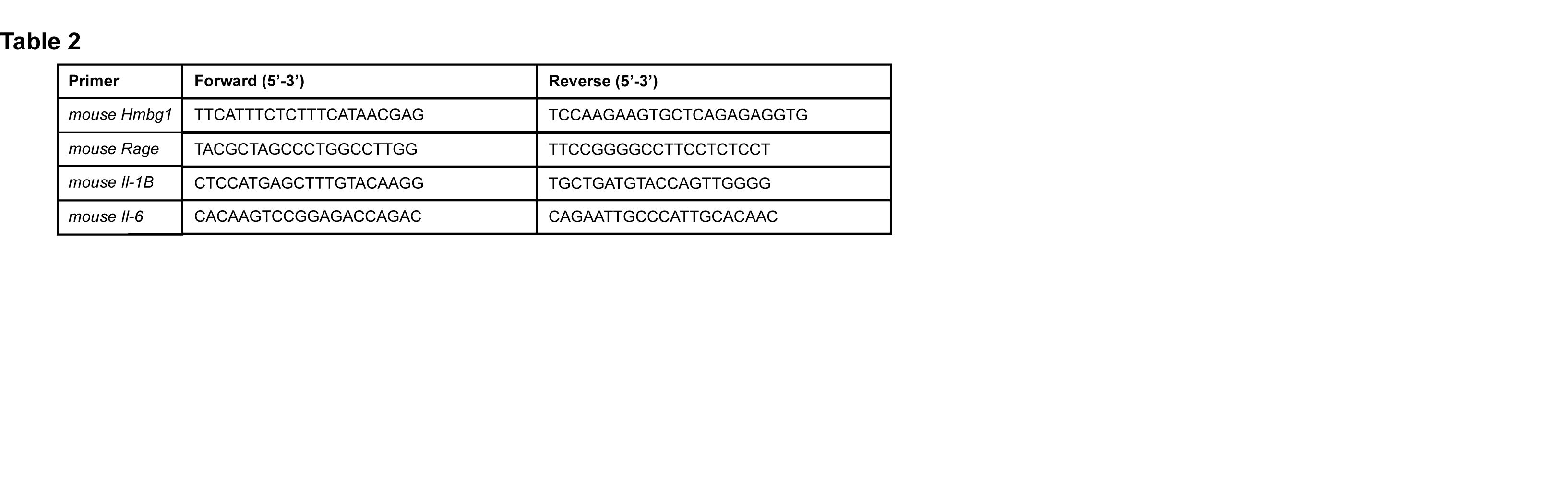
